# Are “early birds” bolder? Early daily activity is not correlated with risk-taking behaviour in a major invasive species

**DOI:** 10.1101/864611

**Authors:** Valerio Sbragaglia, Thomas Breithaupt

**Affiliations:** Department of Marine Renewable Resources, Marine Science Institute (ICM-CSIC), Barcelona Spain; Department of Biology and Ecology of Fishes, Leibniz-Institute of Freshwater Ecology and Inland Fisheries, Müggelseedamm 310, 12587 Berlin, Germany; Department of Biological and Marine Sciences, University of Hull, Hull HU6 7RX, UK

**Keywords:** crayfish, *Pacifastacus leniusculus*, daily activity rhythms, personality, behavioural syndrome

## Abstract

Daily behavioural rhythms provide ecological advantages with respect to exploitation of food resources and avoidance of predation and recent studies suggested that timing of activity could form a behavioural syndrome with risk-taking behavior. Behavioural syndromes are often displayed by invasive species but the role of activity rhythms in biological invasions is unknown. Here, we investigated whether early nocturnal activity (the relative amount of locomotor activity displayed early in the night) and risk-taking behaviour (i.e. response to a scare stimulus) form a behavioural syndrome in a major invasive species, the signal crayfish (*Pacifastacus leniusculus*). We first characterized daily rhythms of locomotor activity over five days under controlled laboratory conditions and then scored the response to a scare stimulus across two different contexts (neutral and food) two days apart within the following six days. Crayfish displayed overall daily activity rhythms peaking in the first four hours of darkness. Both early nocturnal activity and risk-taking behaviour showed consistent inter-individual differences with repeatability scores of 0.20 and 0.35, respectively. However, the two behavioural traits were not correlated as in a behavioural syndrome. We argue that in contrast to other behavioural syndromes, a link between early nocturnal activity and risk-taking tendency would not be evolutionary stable as it dramatically increases predictability and therefore predatory pressure to individuals. We suggest that daily activity rhythms and risk-taking behavior can be important traits in understanding the adaptations underlying biological invasions or other processes of contemporary evolution.

## INTRODUCTION

The behaviour and physiology of almost all animal taxa is characterized by daily activity rhythms (Dunlap and others 2004). Daily activity rhythms are usually synchronized to the day-night cycle, but other environmental or social factors can also play a role (Castillo-Ruiz and others 2012; Hut and others 2012). Daily behavioural rhythms provide ecological advantages with respect to exploitation of food resources and avoidance of predation (Helm and others 2017; Kronfeld-Schor and others 2013; Kronfeld-Schor and Dayan 2003). For example, in the desert golden spiny mouse (*Acomys russatus*) individuals that arrive earlier to a foraging patch gain more food than individuals that arrive later (Levy and others 2012). In zebrafish (*Danio rerio*), individuals displaying higher locomotion early in the activity period were also more risk-taking than individuals that were less active (Tudorache and others 2018). An increase of risk-taking behaviour is usually associated with an increased access to food resources and a decrease of survival towards predators (Lima and Dill 1990; Smith and Blumstein 2008). Therefore, the amount of locomotion displayed early in the activity period could be a fitness-related trait similar to chronotypes (Helm and others 2017).

Chronotypes are consistent inter-individual differences in the onset of activity with respect to a reference time point (Helm and others 2017), and they can have fitness consequences. For example, a recent study suggests that razorfish (*Xyrichtys novacula*) with early-active chronotypes could experience higher mortality from fisheries than late chronotypes (Martorell-Barceló and others 2018). However, daily activity rhythms may not only differ in the onset (i.e. chronotypes) but also in the relative amount of activity displayed at particular time periods. A recent study on zebrafish (*Danio rerio*) showed consistent inter-group differences in the relative amount of locomotion displayed early in the activity period (Sbragaglia and others 2019). Interestingly, elevated locomotion displayed early in the activity period was found to be linked to an increase of risk-taking behaviour in an evolutionary context (Sbragaglia and others 2019), suggesting an adaptive value for such combination of traits.

Animal behaviour plays a paramount role in biological invasions (Holway and Suarez 1999). Personality traits (i.e. consistent inter-individual differences in behaviour; Carere and Maestripieri 2013) such as risk-taking behaviour can determine the success or failure at different stages of biological invasions (Canestrelli and others 2016; Carere and Gherardi 2013; Chapple and others 2012; Juette and others 2015). By contrast, the role of daily activity rhythms in biological invasions is unknown. A key to the understanding of the process of biological invasion could be the finding that successful invaders often display correlation among a suite of consistent behavioural traits (i.e. behavioural syndrome; Chapple and others 2012; Sih and others 2004). For example, a behavioural syndrome between aggression and foraging activity is suggested to enhance the ability of the signal crayfish (*Pacifastasus leniusculus*), a major invasive species, to attain high densities and a negative impact on invaded communities (Pintor and others 2009).

Here, we test the hypothesis that early daily locomotor activity and risk-taking behaviour form a behavioural syndrome in *P. leniusculus*, a commercially harvested crayfish which is one of the most invasive species in Asia and Europe (Gherardi 2017; Larson and others 2012; Lodge and others 2000). We predict that early daily locomotor activity and risk-taking behaviour are consistent behavioural traits and that these traits are positively correlated.

## MATERIALS AND METHODS

### Sampling and acclimation

Crayfish were sourced from a commercial supplier (Flowers Farm Lakes, Dorchester, UK). Prior to experiments crayfish were acclimated in communal tanks (60×45×25 cm) in sex-segregated groups of 10-15 individuals. Crayfish were fed a mixed diet of prawns, peas and carrots. The light-darkness cycle was 12-12 hours and water temperature was kept at 15±1 °C.

We used a total of 24 male crayfish (cephalothorax length: 36.0 ± 2.7 mm). We conducted a total of three experimental trials from May to July. We ran each trial with 8 randomly-selected crayfish in an experimental room where light (light-darkness cycle of 12-12 hours) and water temperature (l5±1 C) were under control. Light was provided by fluorescent lamps at an intensity of 0.25 Klux. During the darkness period dim red light (0.01 Klux) was provided by monochromatic LEDs to allow video recording.

The experimental setup consisted of 8 separated glass aquaria (45×30×25 cm). We equipped each tank with a recirculation pump (provided with a filter). We glued white opaque tissue at the bottom of the tank to create a suitable surface for the crayfish to walk on and the appropriate background to apply video imaging analysis (see below). We placed a shelter (12 cm long) in each tank, cutting transversally a PVC pipe of a diameter of 8 cm. We covered the sides of each tank with white fabric to eliminate any visual disturbance from neighboring tanks and we placed the tanks behind light proof curtains to eliminate any kind of light contamination.

### Experimental trials

We left the crayfish undisturbed during the first stage of the experiment recording a time-lapse video (10 s frequency) that we used to track daily rhythms of locomotor activity (see below). Next, during the second stage we exposed crayfish to a scare stimulus in two different contexts (neutral and food contexts) to evaluate their level of risk-taking behaviour. One of the most important anti-predatory behaviours of crayfish include reduced movement (Stein and Magnuson 1976), therefore we considered the amount of locomotor activity performed 30 min after the scare stimulus as a measure of risk taking behavior (i.e. the less the locomotor activity the less risk-taking behavior). We applied the scare stimulus two days apart within six days moving a plastic black board (15×15 cm) forward and back two times over the surface (see video S1). During the neutral context nothing was changed in the experimental tank, while during the food context a small piece of prawn was added after light-off. The scare stimulus was applied within two hours after light-on only when the crayfish was moving or approaching the food (displacement for more than 1 body length), otherwise the stimulus was not applied and the behavior scored with a 0.

### Behavioral tracking

We quantified locomotor activity using automated video image analysis. Four USB webcams (1 webcam for 2 aquaria) were placed behind light proof curtains on the top of the aquaria. We managed frame acquisition using open source camera security software (ispy: http://www.ispyconnect.com). The automated video image analysis was performed in Matlab 7.1 (The MathWorks, Natick, USA), adapting a compilation of a script with the image processing toolbox previously used in several studies with decapods crustaceans (Sbragaglia and others 2013a; Sbragaglia and others 2015a; Sbragaglia and others 2015b). The final output was a time series of the movement of each crayfish (cm) at a frequency of 10 s.

### Statistical analysis

Chi-square periodogram (Sokolove and Bushell 1978) was used to scan for periodicity and the percentage of variance (%V) was reported as a measure of rhythms’ robustness (Refinetti 2006). Early nocturnal activity was calculated for each of the five days as the proportion between the percent of locomotor activity within four hours after light-off in relation to the percent of time (more details and examples are provided in Figure 1). This provides a quantitative measure of activity within a defined 4-hour time period respect to the overall activity (see also; Sbragaglia and others 2013b).

**Figure 1.**
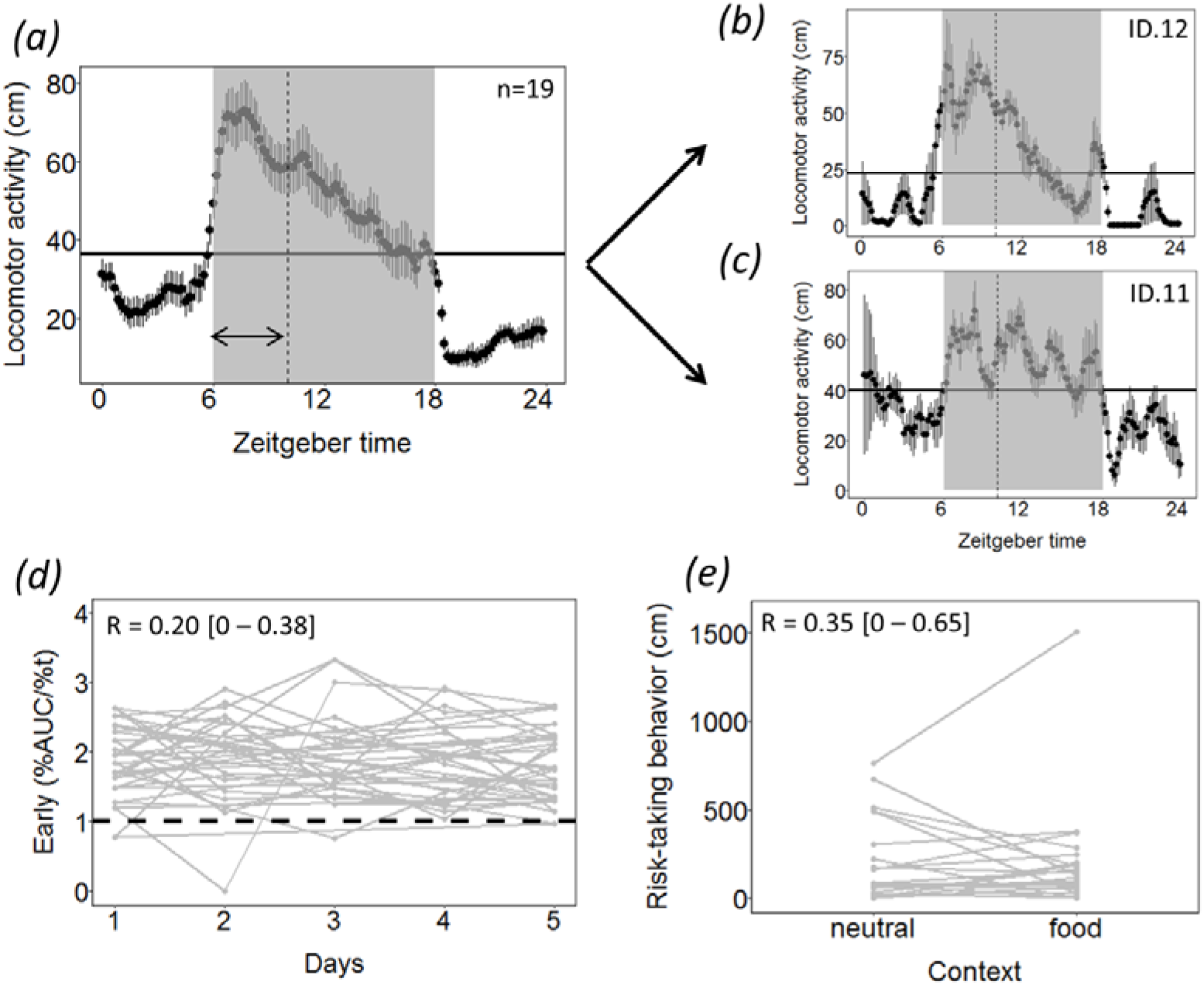
Daily activity rhythms and risk-taking behaviour. (a): Average daily locomotor activity is presented for all individuals together with standard errors (vertical bars) and the midline estimating statistic of rhythm (i.e. mesor, the central tendency of locomotor activity; horizontal line). The vertical dashed line and the double-ended arrow represent the first four hours of darkness where we calculated the proportion between the percent of locomotor activity in relation to the percent of time (%AUC/%t). The grey areas represent darkness hours. (b-c): two representative examples of daily activity rhythms, ID12 (early %AUC/%t = 2.58) and ID11 (early %AUC/%t = 1.36). (d-e): Repeatability of early activity across five days (n = 19) and risk-taking behaviour across contexts (n=24). The horizontal dashed line represents a threshold and values above 1 indicate that locomotor activity is mainly displayed during the first four hours of darkness.

Daily values of early nocturnal activity and the scores for risk-taking behaviour in each context were transformed by finding the exponent (lambda), which made the values of the response variable as normally distributed as possible with a simple power transformation. Repeatability was calculated across time (5 days) for early nocturnal activity and across contexts (neutral and food) for risk-taking behaviour (Nakagawa and Schielzeth 2010). Model fitting was examined by checking normality of residuals and plotting theoretical quantiles vs. standardized residuals. The correlation among the average early nocturnal activity and risk-taking behaviour was calculated with the non-parametric Kendall’s coefficient of concordance.

Analysis were performed using the software Eltemps (www.el-temps.com) and R 3.3.1 (https://www.R-project.org/) with the following additional packages: rcompanion” (https://CRAN.R-project.org/package=rcompanion), “rptR” (Stoffel and others 2017). In all cases, we used a 95% confidence interval.

## RESULTS

Locomotor activity of crayfish during the first stage of the experiment indicated robust daily activity rhythms (mean robustness ± standard deviation = 46±12 %; n = 19; five crayfish did not show significant periodicity in their activity rhythms) with more activity concentrated during darkness (72±11%; n = 19; Fig. 1a). Early nocturnal activity ranged from 1.26 to 2.58 (i.e. the crayfish displayed a 2.58 times more locomotor activity within 4-hours after light-off with respect to overall activity; fig. 1b). Early nocturnal activity was repeatable over time (R = 0.20 [0.01 − 0.38]; p < 0.05; n = 19; Fig. 1d).

During the 30 min after the scaring stimuli, the distance covered by crayfish (risk-taking behavior) ranged from 0 to 760 cm in the neutral context, while it ranged from 0 to 1505 cm in the food context. Two crayfish remained in the shelter (one in the neutral context and one in the food context) during the selected time period (two hours within light-off) in which we applied the stimuli. We scored this behaviour with a 0 because our interpretation is that remaining in the shelter implies a low level of risk-taking behaviour (Jurcak and others 2016). Risk-taking behaviour was significantly repeatable across contexts (R = 0.35 [0.01 − 0.65]; p < 0.05; n = 24; Fig. 1e). Finally, early nocturnal activity did not show significant correlations with risk taking behaviour both in the neutral (r_τ_= −0.03; *p* = 0.860; Fig. 2) and food context (r_τ_= 0.22; *p* = 0.183; fig. 2). Finally, early nocturnal activity was not correlated with the central tendency of locomotor activity (mesor; r_τ_= −0.04; *p* = 0.806).

**Figure 2.**
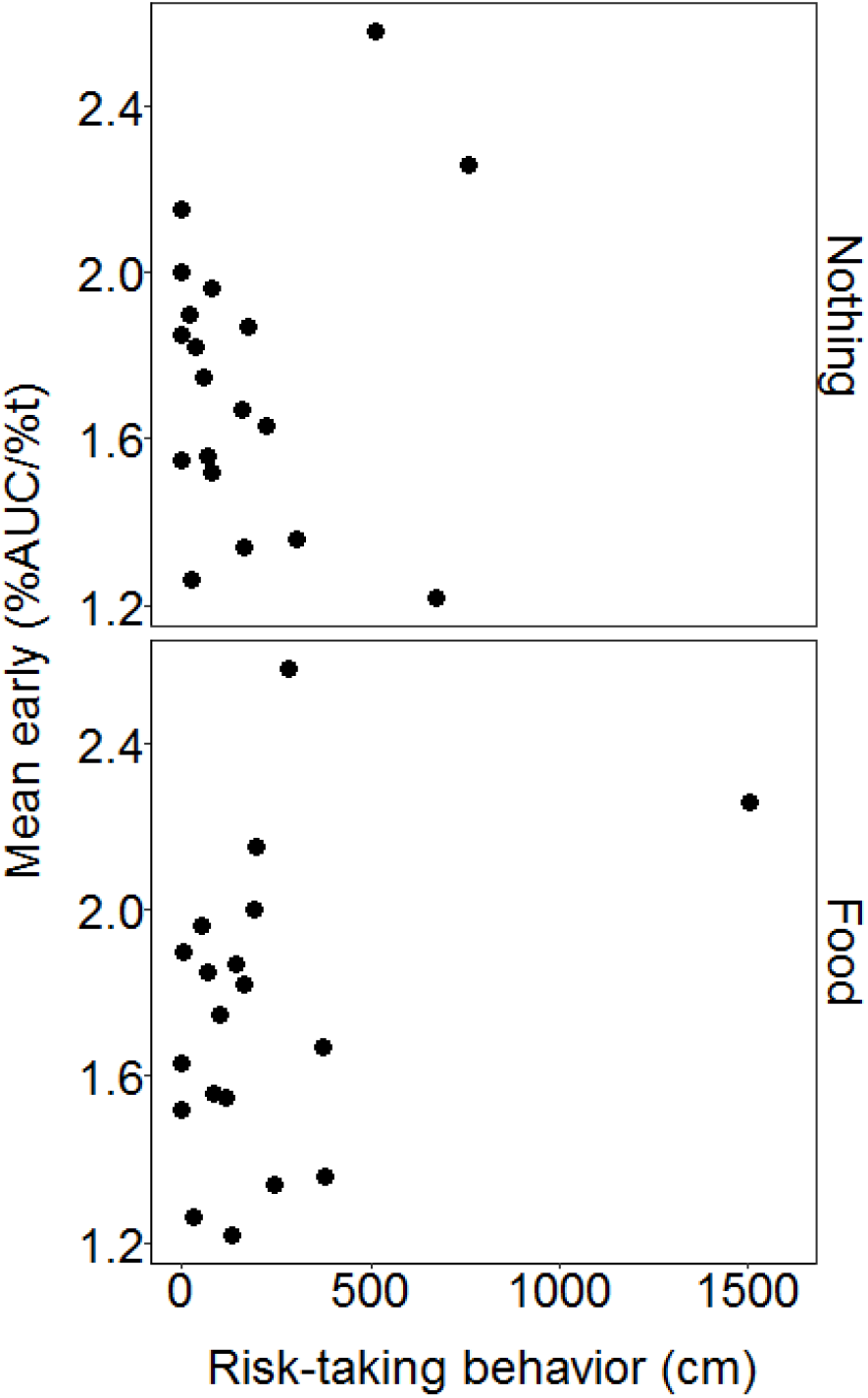
Plot of risk-taking behaviour measured in two different contexts (Nothing and Food) against mean early-active phenotypes (n=19).

## DISCUSSION

The importance of consistent inter-individual differences in the timing of activity has recently been highlighted with respect to the onset of activity (“chronotypes”; Helm and others 2017). In our study we provide evidence that the relative amount of locomotion displayed early in the activity period can also show consistent inter-individual differences. Daily locomotor activity rhythms have a high degree of plasticity. For example, temporal niche switching (i.e. shifts in the timing of daily locomotor activity) has been suggested as a strategy to maximize survival in response to environmental changes both in endotherms and ectotherms (Chiesa and others 2010; Hut and others 2012; van der Vinne and others 2019). Our results suggest that consistent inter-individual differences could be an additional factor in shaping the ecological significance of biological rhythms. Indeed, behavioural ecologists have already recognized that both consistent inter-individual differences (i.e. personality) and individual plasticity must be considered as complementary aspects of individual phenotypes (Dingemanse and others 2010). It must be noted that the early nocturnal activity measured here does not correlate to the mean level of overall activity (i.e. the central tendency of locomotor activity; see Fig. 1a-c). Therefore, the relative amount of locomotion displayed early in the activity period is a trait that is not correlated to overall activity.

Our results provide the first in-depth characterization of daily locomotor activity rhythms of *P. leniusculus*, showing an overall peak of activity in the first hours of darkness (early nocturnal activity) as well as consistency of inter-individual variation. Previous studies already showed that *P. leniusculus* is a nocturnal species (Edmonds and others 2011; Nyström 2005; Thomas and others 2016), but a clear daily activity pattern was never recorded for this major invasive species. The evolution of early nocturnal activity could possibly be related to foraging opportunities as demonstrated in other nocturnal decapod crustaceans such as the Norway lobster, *Nephrops norvegicus* (e.g. Aguzzi and others 2015; Katoh and others 2013; Sbragaglia and others 2013b). Interestingly, we showed that the consistent inter-individual differences in early nocturnal activity account for 20% of the phenotypic variance across time. Similarly, consistent inter-individual differences in risk-taking behaviour account for 35% of the phenotypic variance across contexts. The degree of consistency corresponds to personality traits of other species in a wide range of animal taxa (Bell and others 2009).

The main hypothesis of our study must be rejected because we did not find a behavioural syndrome between early nocturnal activity and risk-taking behaviour. Crayfish with a higher early nocturnal activity could have advantages in finding and exploiting food patches; however, higher activity could increase exposure to predators (Lima and Dill 1990). The major predators of adult crayfish are large predatory fish, wading birds and mammals such as raccoons and otters (Jones and others 2016; Jurcak and others 2016). Important anti-predatory behaviours of crayfish include reduced movement (Stein and Magnuson 1976) or seeking cover (Jurcak and others 2016), and crayfish exposed to fish predators spent less time out of the refuge at night than individuals not exposed to predators (Nyström 2005). The trade-off between anti-predatory behaviour and foraging could explain the adaptive value of the lack of a behavioural syndrome between early nocturnal activity and risk-taking behaviour, suggesting that a common architecture (e.g. pleiotropic genes or common neural pathways) between these traits is unlikely. However, previous results on zebrafish indicated an adaptive value of a behavioural syndrome between these two traits (Sbragaglia and others 2019; Tudorache and others 2018). Decapod crustacean are poorly represented in the studies on animal personality and behavioural syndromes in comparison to fishes, birds and mammals (Gherardi and others 2015). Therefore, we lack comparative studies to understand the general ecological significance of such behavioral syndrome in nocturnal crustaceans.

Biological invasions are often characterized by contemporary evolution (Colautti and Lau 2015; Reisinger and others 2017). For example, common gardens experiments with the rusty crayfish (*Orconectes rusticus*) indicated trait divergence between native and invaded range populations (Sargent and Lodge 2014). In particular, individuals from the invaded populations had faster growth rates and higher survival than individuals from the native range (Sargent and Lodge 2014). Similarly, the signal crayfish (P. *leniusculus*) show changes in population characteristics at the edge of its invasive range in Europe (Hudina and others 2012). In particular, the population at the invasion front was characterized by a male-biased sex ratio and an increase in relative claw size (Hudina and others 2012). Therefore, our results could be also explained according to an evolutionary scenario associated to the invasion process (Chapple and others 2012). Individuals used in this study were obtained from Dorset (Southern UK) where the presence of *P. leniusculus* was already reported more than 40 years ago (Guan and Wiles 1997; Holdich and others 1999). A behavioural syndrome between early nocturnal activity and risk-taking behaviour could have an adaptive advantage during the first stages of invasion, but may lose its adaptive value once crayfish populations are well established. For example, *P. leniusculus* collected at the invasion front in Croatian rivers showed lower level of intra-specific aggression than individuals from the core population, suggesting that a decrease in aggression is maladaptive once the population is established (Hudina and others 2015). Indeed, the sequential stages of invasion could have different selection landscapes that may erode behavioural syndromes (Chapple and others 2012).

A recent study, simulating fisheries-induced evolution, showed that life-history traits, early daily activity and risk-taking behaviour are under selection in population where smaller individuals are selectively harvested (Sbragaglia and others 2019). Although evolutionary adaptations in response to fisheries (Heino and others 2015; Uusi-Heikkilä and others 2015) and biological invasions (Colautti and Lau 2015; Suarez and Tsutsui 2008) are triggering interest among researchers, possible common patterns have seldom been explored. Future studies could benefit from integrating such fields in order to understand the underpinning mechanisms driving contemporary evolution of behavioral syndromes.

## Ethical statement

Experiments were conducted at the University of Hull and approved by the Ethics committee of the Department of Biological Sciences (approval number U035).

## Data accessibility

Data are available as supplementary material together with the R code.

## Authors’ contribution

TB and VS conceived the idea; VS run the experiments, analyzed the data and wrote the manuscript with inputs of TB.

## Acknowledgments

We are thankful to Claudio Carere, Domino Joice, Cristiano Bertolucci and Kate Laskowski for scientific discussion on some of the results presented here. We are also grateful to Jose A. Garcia for his help during animal tracking, Victor Swetez for helping to build the experimental set up and Sonia Jennings for animals’ husbandry.

## REFERENCES

Aguzzi, J.; Sbragaglia, V.; Tecchio, S.; Navarro, J.; Company, J.B. Rhythmic behaviour of marine benthopelagic species and the synchronous dynamics of benthic communities. Deep Sea Res (I Oceanogr Res Pap). 95:1–11; 2015

Bell, A.M.; Hankison, S.J.; Laskowski, K.L. The repeatability of behaviour: a meta-analysis. Anim Behav. 77:771–783; 2009

Canestrelli, D.; Bisconti, R.; Carere, C. Bolder Takes All? The Behavioral Dimension of Biogeography. Trends Ecol Evol. 31:35–43; 2016

Carere, C.; Gherardi, F. Animal personalities matter for biological invasions. Trends Ecol Evol. 28:5–6; 2013

Carere, C.; Maestripieri, D. Animal personalities: behavior, physiology, and evolution: University of Chicago Press; 2013

Castillo-Ruiz, A.; Paul, M.J.; Schwartz, W.J. In search of a temporal niche: social interactions. Prog Brain Res. 199:267–280; 2012

Colautti, R.I.; Lau, J.A. Contemporary evolution during invasion: evidence for differentiation, natural selection, and local adaptation. Mol Ecol. 24:1999–2017; 2015

Chapple, D.G.; Simmonds, S.M.; Wong, B.B.M. Can behavioral and personality traits influence the success of unintentional species introductions? Trends Ecol Evol. 27:57–64; 2012

Chiesa, J.J.; Aguzzi, J.; García, J.A.; Sardà, F.; de la Iglesia, H.O. Light intensity determines temporal niche switching of behavioral activity in deep-water *Nephrops norvegicus* (Crustacea: Decapoda). J Biol Rhythms. 25:277–287; 2010

Dingemanse, N.J.; Kazem, A.J.N.; Réale, D.; Wright, J. Behavioural reaction norms: animal personality meets individual plasticity. Trends Ecol Evol. 25:81–89; 2010

Dunlap, J.C.; Loros, J.J.; DeCoursey, P.J. Chronobiology: biological timekeeping. Sunderland Massachusetts: Sinauer Associates; 2004

Edmonds, N.; Riley, W.; Maxwell, D. Predation by *Pacifastacus leniusculus* on the intra-gravel embryos and emerging fry of *Salmo salar*. Fish Manage Ecol. 18:521–524; 2011

Gherardi, F. Crayfish in Europe as alien species: Routledge; 2017

Gherardi, F.; Aquiloni, L.; Tricarico, E. Behavioral plasticity, behavioral syndromes and animal personality in crustacean decapods: An imperfect map is better than no map. Curr Zool. 58:567–579; 2015

Guan, R.Z.; Wiles, P.R. Ecological impact of introduced crayfish on benthic fishes in a British lowland river. Conserv Biol. 11:641–647; 1997

Heino, M.; Díaz Pauli, B.; Dieckmann, U. Fisheries-induced evolution. Annu Rev Ecol, Evol Syst. 46:461–480; 2015

Helm, B.; Visser, M.E.; Schwartz, W.; Kronfeld-Schor, N.; Gerkema, M.; Piersma, T.; Bloch, G. Two sides of a coin: ecological and chronobiological perspectives of timing in the wild. Phil Trans R Soc B. 372:20160246; 2017

Holdich, D.M.; David Rogers, W.; Reynolds, J.D. Native and alien crayfish in the British Isles. Crustacean Issues. 11:221–236; 1999

Holway, D.A.; Suarez, A.V. Animal behavior: an essential component of invasion biology. Trends Ecol Evol. 14:328–330; 1999

Hudina, S.; Hock, K.; Žganec, K.; Lucić, A. Changes in population characteristics and structure of the signal crayfish at the edge of its invasive range in a European river. Annales de Limnologie - International Journal of Limnology. 48:3–11; 2012

Hudina, S.; Žganec, K.; Hock, K. Differences in aggressive behaviour along the expanding range of an invasive crayfish: an important component of invasion dynamics. Biol Invasions. 17:3101–3112; 2015

Hut, R.A.; Kronfeld-Schor, N.; van der Vinne, V.; De la Iglesia, H. In search of a temporal niche: environmental factors. Prog Brain Res. 199:281–304; 2012

Jones, E.W.; Jackson, M.C.; Grey, J. Environmental drivers for population success: Population biology, population and community dynamics. Biology and Ecology of Crayfish: 251–286; 2016

Juette, T.; Cucherousset, J.; Cote, J. Animal personality and the ecological impacts of freshwater non-native species. Curr Zool. 60:417–427; 2015

Jurcak, A.M.; Lahman, S.E.; Wofford, S.J.; Moore, P.A. Behavior of crayfish. Biology and Ecology of Crayfish; Longshaw, M, Stebbing, B, Eds:117–131; 2016

Katoh, E.; Sbragaglia, V.; Aguzzi, J.; Breithaupt, T. Sensory biology and behaviour of *Nephrops norvegicus*. in: Johnson M.L., Johnson M.P., eds. The ecology and biology of Nephrops norvegicus: Acedemic Press; 2013

Kronfeld-Schor, N.; Bloch, G.; Schwartz, W.J. Animal clocks: when science meets nature. Proc R Soc Lond B. 280:20131354; 2013

Kronfeld-Schor, N.; Dayan, T. Partitioning of time as an ecological resource. Annu Rev Ecol, Evol Syst. 34:153–181; 2003

Larson, E.R.; Abbott, C.L.; Usio, N.; Azuma, N.; Wood, K.A.; Herborg, L.M.; Olden, J.D. The signal crayfish is not a single species: cryptic diversity and invasions in the Pacific Northwest range of *Pacifastacus leniusculus*. Freshwat Biol. 57:1823–1838; 2012

Levy, O.; Dayan, T.; Rotics, S.; Kronfeld-Schor, N. Foraging sequence, energy intake and torpor: an individual-based field study of energy balancing in desert golden spiny mice. Ecol Lett. 15:1240–1248; 2012

Lima, S.L.; Dill, L.M. Behavioral decisions made under the risk of predation: a review and prospectus. Can J Zool. 68:619–640; 1990

Lodge, D.M.; Taylor, C.A.; Holdich, D.M.; Skurdal, J. Nonindigenous crayfishes threaten North American freshwater biodiversity: lessons from Europe. Fisheries. 25:7– 20;2000

Martorell-Barceló, M.; Campos-Candela, A.; Alós, J. Fitness consequences of fish circadian behavioural variation in exploited marine environments. PeerJ. 6:e4814;2018

Nakagawa, S.; Schielzeth, H. Repeatability for Gaussian and non-Gaussian data: a practical guide for biologists. Biol Rev Camb Philos Soc. 85:935–956; 2010

Nyström, P. Non-lethal predator effects on the performance of a native and an exotic crayfish species. Freshwat Biol. 50:1938–1949; 2005

Pintor, L.M.; Sih, A.; Kerby, J.L. Behavioral correlations provide a mechanism for explaining high invader densities and increased impacts on native prey. Ecology. 90:581–587; 2009

Refinetti, R. Circadian physiology (2nd ed). CRC Press: Boca Raton, Fla; 2006

Reisinger, L.S.; Elgin, A.K.; Towle, K.M.; Chan, D.J.; Lodge, D.M. The influence of evolution and plasticity on the behavior of an invasive crayfish. Biol Invasions. 19:815–830; 2017

Sargent, L.W.; Lodge, D.M. Evolution of invasive traits in nonindigenous species: increased survival and faster growth in invasive populations of rusty crayfish (*Orconectes rusticus*). Evol Appl. 7:949–961; 2014

Sbragaglia, V.; Aguzzi, J.; García, J.; Sarriá, D.; Gomariz, S.; Costa, C.; Menesatti, P.; Vilaró, M.; Manuel, A.; Sardà, F. An automated multi-flume actograph for the study of behavioral rhythms of burrowing organisms. J Exp Mar Biol Ecol. 446:177–185; 2013a

Sbragaglia, V.; Aguzzi, J.; Garcia, J.A.; Chiesa, J.J.; Angelini, C.; Sardà, F. Dusk but not dawn burrow emergence rhythms of *Nephrops norvegicus* (Crustacea: Decapoda). Sci Mar. 77:641–647; 2013b

Sbragaglia, V.; García, J.; Chiesa, J.; Aguzzi, J. Effect of simulated tidal currents on the burrow emergence rhythms of the Norway lobster (*Nephrops norvegicus*). Mar Biol. 162:2007–2016; 2015a

Sbragaglia, V.; Lamanna, F.; M. Mat, A.; Rotllant, G.; Joly, S.; Ketmaier, V.; de la Iglesia, H.O.; Aguzzi, J. Identification, characterization, and diel pattern of expression of canonical clock genes in *Nephrops norvegicus* (Crustacea: Decapoda) eyestalk. PLoS One. 10:e0141893; 2015b

Sbragaglia, V.; López-Olmeda, J.F.; Frigato, E.; Bertolucci, C.; Arlinghaus, R. Fisheries-induced evolution of the circadian system and collective personality traits. bioRxiv:622043; 2019

Sih, A.; Bell, A.; Johnson, J.C. Behavioral syndromes: an ecological and evolutionary overview. Trends Ecol Evol. 19:372–378; 2004

Smith, B.R.; Blumstein, D.T. Fitness consequences of personality: a meta-analysis. Behav Ecol. 19:448–455; 2008

Sokolove, P.G.; Bushell, W.N. The chi square periodogram: its utility for analysis of circadian rhythms. J Theor Biol. 72:131–160; 1978

Stein, R.A.; Magnuson, J.J. Behavioral response of crayfish to a fish predator. Ecology. 57:751–761; 1976

Suarez, A.V.; Tsutsui, N.D. The evolutionary consequences of biological invasions. Mol Ecol. 17:351–360; 2008

Thomas, J.R.; James, J.; Newman, R.C.; Riley, W.D.; Griffiths, S.W.; Cable, J. The impact of streetlights on an aquatic invasive species: Artificial light at night alters signal crayfish behaviour. Appl Anim Behav Sci. 176:143–149; 2016

Tudorache, C.; Slabbekoorn, H.; Robbers, Y.; Hin, E.; Meijer, J.H.; Spaink, H.P.; Schaaf, M.J.M. Biological clock function is linked to proactive and reactive personality types. BMC Biol. 16:148; 2018

Uusi-Heikkilä, S.; Whiteley, A.R.; Kuparinen, A.; Matsumura, S.; Venturelli, P.A.; Wolter, C.; Slate, J.; Primmer, C.R.; Meinelt, T.; Killen, S.S.; Bierbach, D.; Polverino, G.; Ludwig, A.; Arlinghaus, R. The evolutionary legacy of size-selective harvesting extends from genes to populations. Evol Appl. 8:597–620; 2015

van der Vinne, V.; Tachinardi, P.; Riede, S.J.; Akkerman, J.; Scheepe, J.; Daan, S.; Hut, R.A. Maximising survival by shifting the daily timing of activity. Ecol Lett. 22:2097–2102; 2019

